# WHISKiT Physics: A three-dimensional mechanical model of the rat vibrissal array

**DOI:** 10.1101/862839

**Authors:** Nadina O. Zweifel, Nicholas E. Bush, Ian Abraham, Todd D. Murphey, Mitra J.Z. Hartmann

**Author notes:** **Corresponding Author**: Mitra J.Z. Hartmann, Mechanical Engineering Department, Northwestern University, 2145 Sheridan Rd, Evanston, IL 60208.

## Abstract

Rodents tactually explore the environment using ~62 whiskers (vibrissae), regularly arranged in arrays on both sides of the face. The rat vibrissal system is one of the most commonly used models to study how the brain encodes and processes somatosensory information. To date, however, researchers have been unable to quantify the mechanosensory input at the base of each whisker, because the field lacks accurate models of three-dimensional whisker dynamics. To close this gap, we developed *WHISKiT Physics*, a simulation framework that incorporates realistic morphology of the full rat whisker array to predict time-varying mechanical signals for all whiskers. The dynamics of single whiskers were optimized based on experimental data, and then validated against free tip oscillations and the dynamic response to collision. The model is then extrapolated to include all whiskers in the array, taking into account each whisker’s individual geometry. Simulations of first mode resonances across the array approximately match previous experimental results and fall well within the range expected from biological variability. Finally, we use *WHISKiT Physics* to simulate mechanical signals across the array during three distinct behavioral conditions: passive whisker stimulation, active whisking against two pegs, and active whisking in a natural environment. The results demonstrate that the simulation system can be used to predict input signals during a variety of behaviors, something that would be difficult or impossible in the biological animal. In all behavioral conditions, interactions between array morphology and individual whisker geometry shape the tactile input to the whisker system.

## 1. Introduction

Across sensory modalities, quantifying and analyzing input signals have played a major role in our understanding of sensory processing and perception. For example, images and sound recordings have been indispensable in developing models of the visual and auditory system, from early processing in the periphery (1–3) to cognitive and cortical functions (4–9). Access to the sensory information acquired by these systems has allowed investigators to test and explore models of receptive field representations (6, 10–14), object recognition and speech perception (8, 15–17), saliency and attention (18–22), and neural processing (7).

Compared to the relative ease with which cameras and microphones can capture features of visual and auditory scenes, measuring the sensory input of the somatosensory system is more challenging and, in many cases, impossible. Therefore, tools to simulate somatosensory input are crucial. they will ultimately enable the field to take approaches, such as analysis of natural scene statistics (23–26), that are well established for vision and audition but have remained largely unexplored for somatosensation. Although some models of sensory input have been developed for the primate hand (27, 28), we still lack a model of the complete mechanosensory input to the rodent vibrissal array, the most widely used animal model in somatosensory research.

Here we describe a novel simulation framework (WHISKiT Physics) that incorporates a three-dimensional (3D) dynamical model of the rat vibrissal array to allow researchers to simulate mechanosensory input during active whisking behavior. The model incorporates the typical shape and curvature of each individual whisker as well as the morphology of the rodent’s face and the arrangement of the whiskers on the mystacial pad. Each whisker can be actuated according to typical equations of motion for whisking (29). We present results for the rat, but the model is easily extended to include the mouse.

The model is based on the Bullet Physics Library (30), an open-source physics engine with increasing contributions to game development, robotic simulation, and reinforcement learning (31). This engine provides our model with reasonable computational efficiency, the ability to model natural environments based on 3D polygon meshes, and visualization of the simulations. Because it permits direct control of whisker motion and simultaneous readout of mechanosensory feedback, WHISKiT enables closed-loop simulations of the entire somatosensory modality in the rat.

We first validated the model of a single whisker against experimental data, and then evaluated its generalizability across all whiskers of the array. Model dynamics were tested against analytical solutions of shock wave magnitude and against experimental data describing shock wave propagation after collision with an object (32). Finally, we use the model to simulate vibrissotactile sensory input in three typical exploratory scenarios in laboratory and in natural environment.

## 2. Results

### 2.1 Optimization of single-whisker dynamics in two dimensions

We began by optimizing the dynamics of a model of a single whisker with known arc length, curvature, and base diameter. As described in *Methods*, the single-whisker model has two free parameters representing Young’s modulus (*θ*_E_) and damping (*θ*_*ζ*_). These parameters were optimized based on experiments in which large, caudal whiskers (α and B1) were attached to a motor and driven with a Gaussian pulse at three different speeds. Error was computed by calculating the median symmetric accuracy of resonance frequency (*f*_*n*_) and logarithmic decrement (δ) between simulation and experiment (*Methods*, eq. 4).

To avoid local minima, we performed a brute force search in the two-dimensional parameter space. The resulting error heatmap (Figure 1A) exhibits a linear relationship between *θ*_*E*_ and *θ*_*ζ*_, reflecting the tradeoff between fitting the resonance frequency and fitting the log-decrement. The minimum is located at *θ*_*E*_ = 5.0 GPa and *θ*_*ζ*_ = 0.33. Note that *θ*_*E*_ and *θ*_*ζ*_ were optimized based on the assumption that they are constant within and across whiskers, so they are expected only to approximate the material properties of real whiskers. Values of Young’s modulus have been reported to fall between 1.3-7.8 GPa (33–37), while the damping ratio has been estimated between 0.05 and 0.28 (33, 34, 38).

**Figure 1.**
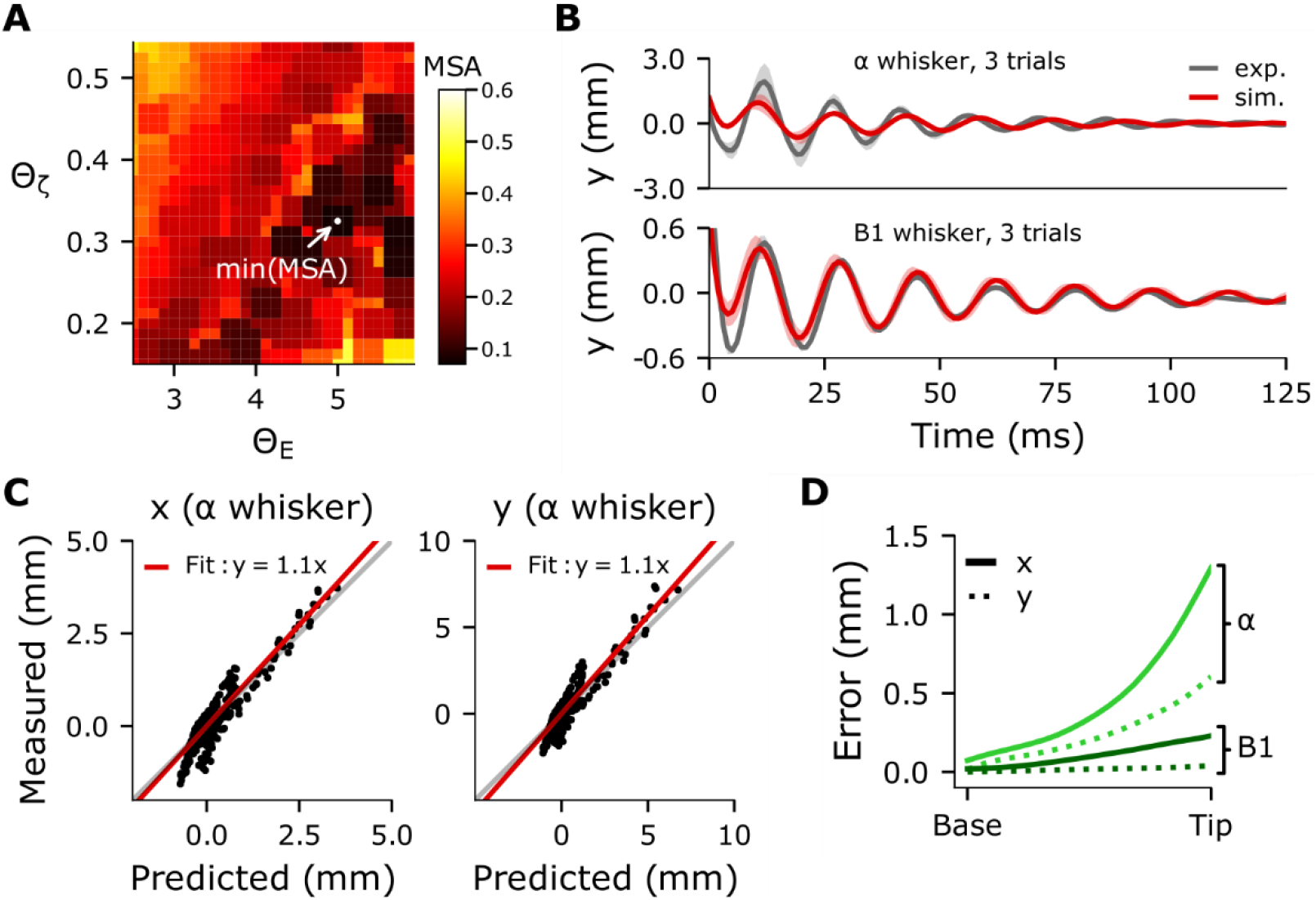
Smooth error surface of parameter optimization with a single whisker yields good match with experimental data. **(A)** Error surface generated by optimizing the parameters *θ*_*ζ*_ and *θ*_*E*_. Color represents error between simulated and experimental data. **(B)** Averaged trajectory of the whisker tip in the y-direction for three trials in which the α and B1 whiskers were driven at the same speed in experiment and simulation. The variability indicated by the standard deviation across the trials occurs because the motor angle is non-deterministic. **(C)** Measured versus predicted trajectory (x- and y- coordinates) of the α whisker tip in the horizontal plane over all trials used for the optimization. **(D)** Displacement error increases as a function of link position (distance from whisker base). Note that the error of the tip, which was used for optimization, is largest; error decreases rapidly to less than 0.3 mm at 50% of the whisker length. Data are shown for the α (green) and B1 (black) whiskers; x- and y- coordinates are shown as dashed and solid lines, respectively.

Given these optimized parameters, the actual dynamic properties measured from simulated trajectories were in good agreement with values observed experimentally (Table 1), and well within observed biological variability. Whiskers with dimensions similar to the α whisker have first-mode resonance frequencies between 59 – 83 Hz; those with dimensions similar to B1, between 50 – 59 Hz (33, 34). The damping ratio calculated from the kinematic behavior of the whisker is smaller than the optimized value for parameter *θ*_*ζ*_. This difference is likely attributable to the assumption that the damping ratio is uniform along the whisker, as previous studies have suggested that the damping ratio decreases from base to tip (32, 33, 38).

**Table 1.** Dynamic measurements of experiment and optimized simulation.

| Variable and units | whisker | Experiment | Simulation | Absolute Error |
| --- | --- | --- | --- | --- |
| $f_n$ (Hz) | $\alpha$ | 67.6 ($\pm 0.72$) | 64.87 ($\pm 1.12$) | 2.73 ($\pm 1.28$) |
| | B1 | 59.38 ($\pm 0.44$) | 58.96 ($\pm 0.68$) | 0.86 ($\pm 0.61$) |
| $\delta$ (dimensionless) | $\alpha$ | 0.32 ($\pm 0.23$) | 0.50 ($\pm 0.4$) | 0.26 ($\pm 0.37$) |
| | B1 | 0.25 ($\pm 0.08$) | 0.23 ($\pm 0.08$) | 0.04 ($\pm 0.06$) |
| $\zeta$ (dimensionless) | $\alpha$ | 0.05 ($\pm 0.04$) | 0.08 ($\pm 0.06$) | 0.04 ($\pm 0.06$) |
| | B1 | 0.04 ($\pm 0.01$) | 0.04 ($\pm 0.01$) | 0.01 ($\pm 0.01$) |

Matching material properties between simulation and experiment does not guarantee that simulated dynamics will accurately match experimental data. Nevertheless, simulated trajectories of the whisker tip closely followed experimentally obtained trajectories (Figure 1B). The high accuracy of the predictions is confirmed by a fit close to the identity line in Figure 1C for the α whisker, with a slope of 1.1 for both x- and y- coordinates. Results for the B1 whisker were similar but had a slightly larger bias in the y direction, with a slope of 1.3. Note that the trajectory error shown in Figure 1B and Figure 1C are the largest observed, because the error increases with the distance to the actuation point (i.e., the whisker base). As illustrated in Figure 1D, the maximum error at the tip is 1.6 mm and decreases rapidly to less than 1 mm at 80% of the whisker length. At 50%, the error is already less than 0.3 mm and nearly zero (< 0.1 mm) at the base.

### 2.2. Resonant frequencies with optimized material parameters generalize across whisker identities

We next quantified how well the model generalized over all whiskers in the array, without reoptimizing any parameters. Using the values *θ*_*E*_ = 5.0 GPa and *θ*_*ζ*_ = 0.33 obtained previously, we quantified model performance against an experimental dataset that characterized the geometry and resonances of 24 whiskers from a single rat (Hartmann et al., 2003; supplementary table 1). Results, shown in Figure 2A, yield a linear fit between predicted and measured resonant frequencies with a slope of 1.0 and an intercept of −31 Hz (solid line). This fit closely matches that obtained from an analytical resonance model, which had a slope of 0.93 and an intercept of 6.4 Hz (33).

**Figure 2.**
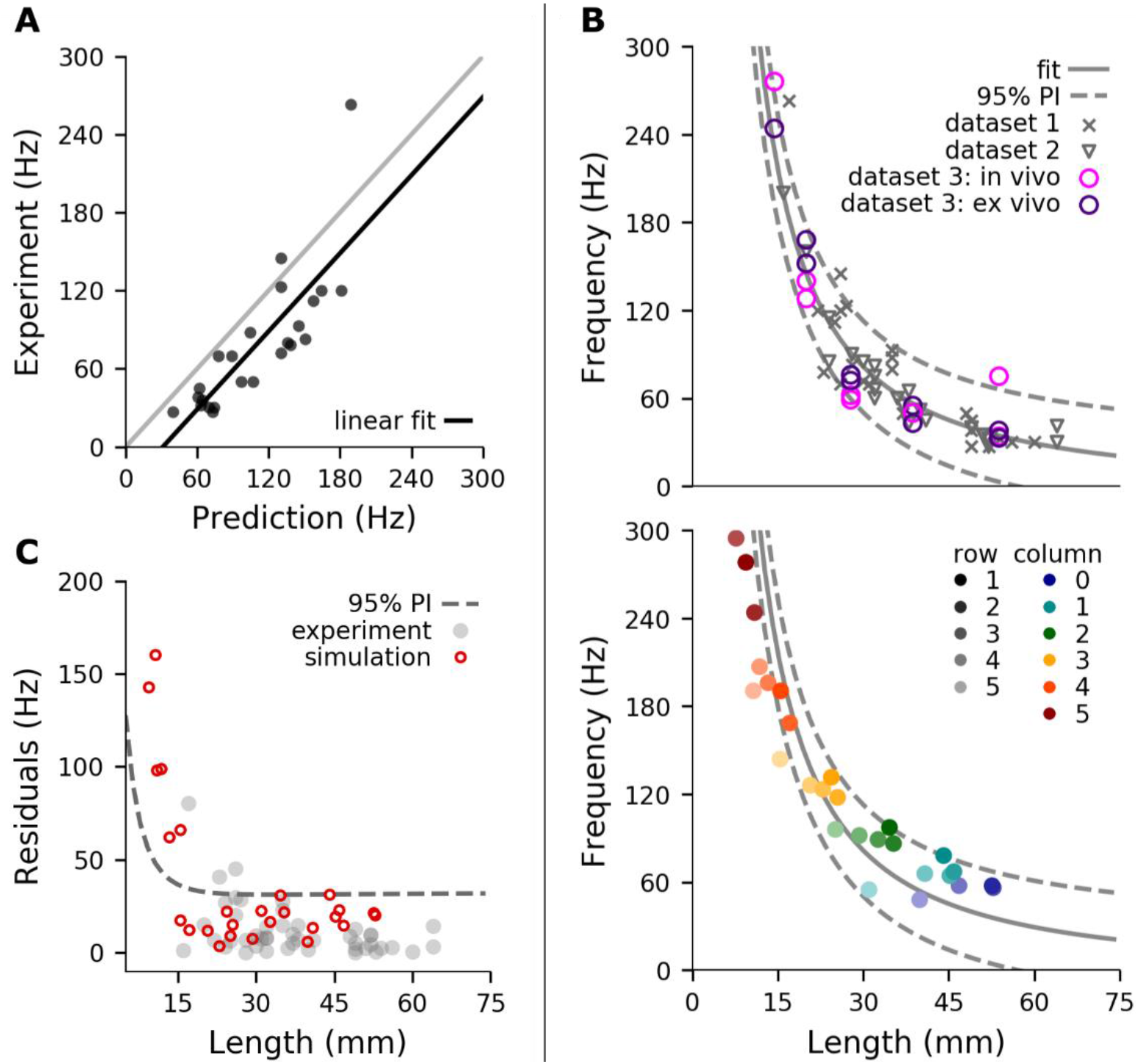
The optimized whisker model generalized across the array. **(A)** Predicted resonance frequencies for dataset 1, obtained from Hartmann et al., 2003 (33). Predictions were obtained from simulations that used the experimentally-measured arc length and base/tip diameter. The gray line represents y = x. **(B)** The relationship between whisker length and resonance frequency obtained from experimental and simulated data. *Upper panel*: Experimentally measured resonance frequencies of the combined dataset including a total of 46 whisker (supplementary table 1). The grey solid line indicates the nonlinear fit (y = 8963 x^(−1.4)^ − 3.8), while the dashed lines indicate the corresponding prediction intervals. *Lower panel*: Simulated resonance frequencies of average whiskers for the right side of the full array (27 whiskers; left side is identical), color coded by whisker identity (row, column). The solid and dashed gray lines are the identical nonlinear fit and prediction intervals shown in the upper panel. **(C)** Residuals of experimental (grey) and simulated (red) resonance frequencies in respect to the nonlinear fit shown in (B top). The prediction interval is identical to the upper bound shown in (B) and is indicated by the dashed line.

Model validation was then extended to incorporate two additional experimental datasets for which only partial geometric and resonance information was available (supplementary table 1). The study of Wolfe et al. (2008) extends the dataset by 22 whiskers (δ, D1, D2, D3, D4) from four different rats (39). Neimark et al. (2003) provides resonance frequencies of 10 whiskers of the C-row (left and right) measured both *in vivo* and *ex vivo* (34). Because lengths for these whiskers were not published, we estimated them from the equations of Belli et al. 2017 (40). All three datasets are plotted in the top panel of Figure 2B. Note that measurement condition (*in vivo* vs. *ex vivo*) is associated with a large spread in the measurements; these differences provide an intuition for the error range associated with experimental data based on the inverse relationship between resonant frequency and whisker length, we fit a negative power function to the three datasets and obtained the model: y = 8963 x^(−1.4)^ − 3.8 with a r^2^ = 0.91 (solid line). The 95% prediction intervals are shown as dashed lines.

Based on this experimental fit, we validated the resonance response across all whiskers of the full array, using average geometry (arc length, base diameter, slope) for each whisker identity as predicted by the models published in Belli et al. 2017 (40). The lower panel of Figure 2B illustrates that the resonance frequencies of the averaged whiskers are well within the prediction intervals obtained from the experimental data in the upper panel (dashed lines). In addition, these resonance frequencies do not significantly differ from the resonance frequencies obtained from simulations using the actual geometry shown in Figure 2A (2-sample t-test: p=0.93).

Plotting the residuals of the regression model in Figure 2B reveals that only whiskers shorter than 15 mm (typically column 5 and higher) fall outside of the prediction intervals (Figure 2C). This error is mainly attributable to the high stiffness and small size of these whiskers, which are challenging for numerical solvers. However, note the large deviation in high frequency measurements and sparsity of data points (Figure 2A, 2B top), which also contribute to the mismatch between experimental data and simulation results.

### 2.3. Dynamic behavior during collision matches previous experimental results for straight and curved whiskers

The results above validate model dynamics during motions of the whisker that do not involve contact. To test the model under conditions of contact and in collision scenarios, we compared it with an analytical model and with experimental data published previously (32). The analytical model of a straight whisker suggests that the mechanical response to a shock event scales with collision velocity, while experimental data indicates that shock wave propagation from tip to base is approximately linear (32).

To replicate the procedures used to test the analytical model of collision (32), we simulated a straight whisker as anchored at its base and rotated it with constant velocity about the vertical axis through the base point until the 19th link (95% of whisker length from the base) collided with a rectangular edge. The collision angle between the whisker and the edge was set to 60° from the horizontal plane, and 45° degrees from the vertical plane to generate mechanical responses in all three dimensions (see schematic in Figure 3A). We simulated collisions at six speeds, for 12 whiskers from four different rows.

**Figure 3.**
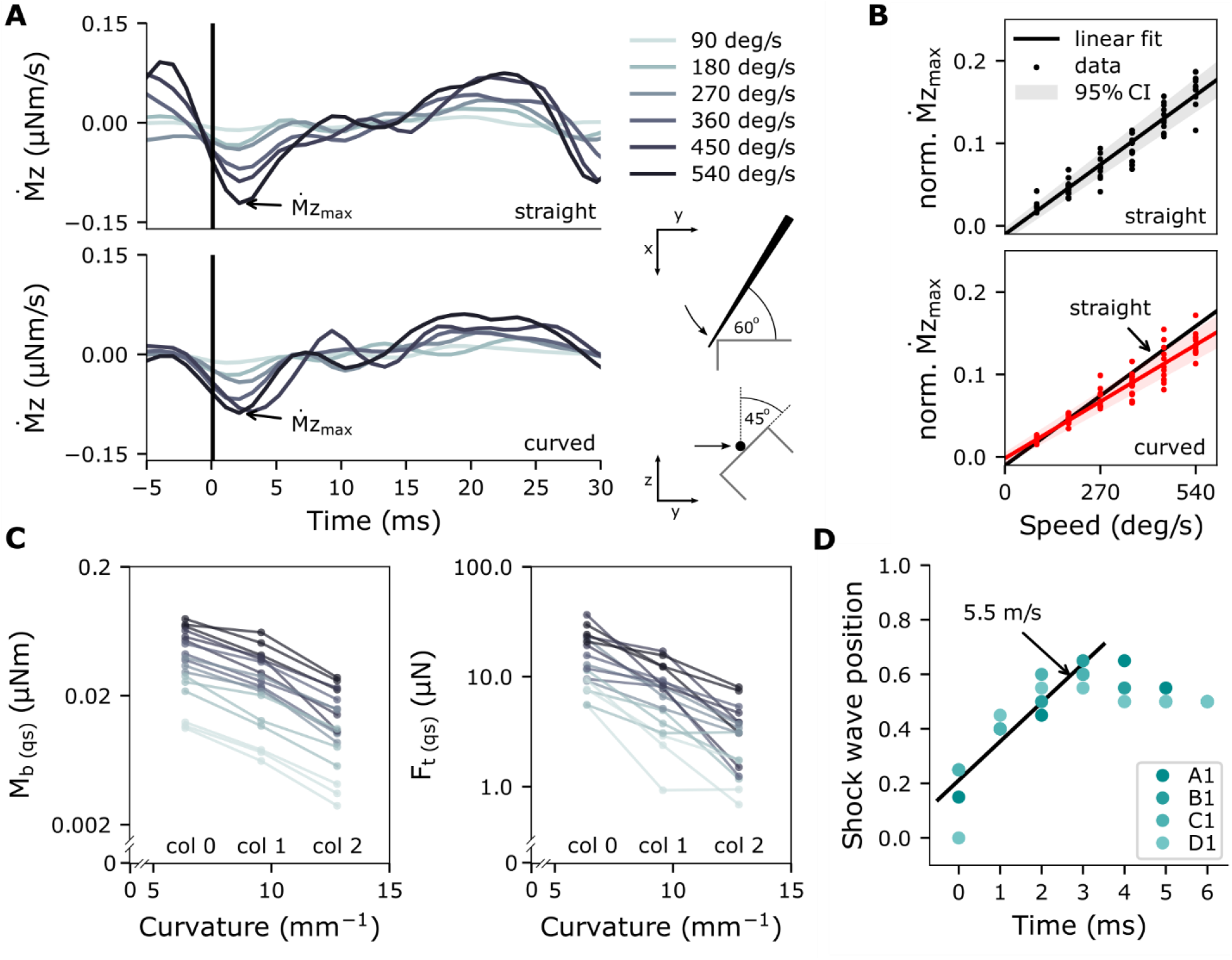
Shock waves resulting from collisions of straight and curved whiskers against a straight edge. **(A)** Mechanical response of a straight and curved whisker in response to a collision, . The derivative of the moment at the base of the α-whisker is shown for six different whisker speeds (90, 180, 270, 360, 450, 540 deg/s; color coded). Trials are aligned at time of impact (black line). **(B)** Relationship between normalized amplitude of shock wave and whisker velocity for 12 straight and 12 curved whiskers (α, A1, A2, B1, β, B1, B2, γ, C1, C2, δ, D1, D2). Linear fit is represented by black solid line (95% confidence interval shaded in grey). Note that the slope of the two regression lines is significantly different between straight and curved whiskers. **(C)** Bending moment M_b_ and transverse force F_t_ at time of maximum shock wave amplitude in respect to curvature of the whisker including linear fit. **(D)** Shock wave position of normalized whisker length for the first 4 rows in the first column of the array. A linear fit over the first 4 ms is shown with solid line. The slope of the linear fit is 5.5 m/s. Color convention is consistent with previous figures.

We then repeated the simulation experiments for a whisker curved according to the model in (41). One of the main advantages of the present numerical model is that intrinsic curvature of the whisker can be included, which is expected to have a significant influence on the mechanical response at the base during collision (42).

Figure 3A shows *Ṁ*_*z*_, the derivative of the moment at the base about the rotation axis for both a straight and curved β whisker, from 5 ms before to 30 ms after collision. In both cases, the first negative peak of *Ṁ*_z_ (shock,*Ṁ*_*z,max*_) occurs ~3 ms after collision, and its magnitude clearly increases with velocity. These results are consistent with analytical solutions of a single straight whisker model (32). We repeated the same simulations for all whiskers in the array and measured *Ṁ*_*z,max*_ for all trials and whiskers. For each whisker, *Ṁ*_*z,max*_ was normalized by its mean value across the six different-speed trials. This normalization allowed us to compare trends across all whiskers of the array. Figure 3B shows normalized *Ṁ*_*z,max*_ value as a function of angular speed at time of collision. *Ṁ*_*z,max*_ correlates with speed, achieving an adjusted r^2^ of 0.92 for both straight and curved whiskers. However, the slopes of the regression lines are significantly different (p<0.001).

The decrease in *Ṁ*_*z,max*_ for curved whiskers can be explained by the negative correlation between curvature and the transverse force (*F_t_*) and between curvature and the bending moment (*M_b_*) at the whisker base, as shown in Figure 3C. Note that *F_t_* and *M_b_* are averaged values of the first 30 ms after collision, representing quasi-static (qs) measures. The slope of the linear regression curve is significant for *M_b_* (r^2^ = 0.307, p < 0.001), as well as *F_t_* (r^2^ = 0.467, p < 0.001). This nearly-linear inverse relationship between the quasi-static mechanical signals at the base and curvature matches predictions from a previous quasistatic numerical model (42).

Figure 3D shows the shock wave position relative to the normalized whisker length within the first 6 ms after collision for 4 whiskers (first 4 rows, first column). A linear regression was performed on the first 4 data points after collision. The slope is significant (r^2^ = 0.90, p < 0.001) and corresponds to a shock wave velocity of 5.5 m/s close to the analytical solution of 5.6 m/s and experimentally measured shock wave velocity of 5.02 m/s (32). Boubenec et al. found the shock wave propagation to be linear within the first 6 ms after collision. We attribute the non-linearities in our model after 3 ms to the reduced spatial resolution of the whisker and increasing stiffness towards the whisker base. We found similar results for the Greek column, which yielded a shock wave velocity of 4.3 m/s (r^2^ = 0.97, p < 0.001).

### 2.4 Damping properties of the follicle considerably impacts the dynamical signals at the whisker base

Having optimized and validated our model based on rigidly anchored whiskers, we next aimed to incorporate elasticity at the whisker base, simulating the insertion of the whisker-follicle complex into compliant skin tissue. For these tests, we used experimental data obtained in the anesthetized animal. We quantified the trajectory of the whisker as it relaxed after a manual deflection delivered with a rigid probe in two directions (43) (see *Methods*).

Five different whisker identities from 11 different animals were used. Contact between probe and whisker occurred at approximately 60% of the whisker length (link 12), but only kinematic data from the most distal link (link 20) during the first 40-50 ms after release from deflection were analyzed. This choice allowed us to avoid any forced motion induced by the deflecting probe. Following the conventions of previous studies (43, 44), the horizontal direction was defined to be parallel to the plane of the intrinsic curvature of the whisker and the vertical direction was perpendicular to that plane.

Preliminary analysis immediately indicated that the trajectory of the whisker tip was considerably more damped than the dynamics of the rigidly anchored whisker used in the previous simulations. Simulations that assumed a rigid follicle generated deflections that were nearly an order of magnitude larger than those observed experimentally (Figure 4A, top panel). We therefore added two identical torsional springs about the y and z axis of the base of the whisker to account for the compliant tissue properties of the follicle. The spring stiffness and damping of these springs were then optimized using randomly selected trials from the B1, B2, and D2 whiskers (*Methods*). After optimization, the trajectory of the whisker more closely resembled the experimental data (Figure 4A, bottom panel).

**Figure 4.**
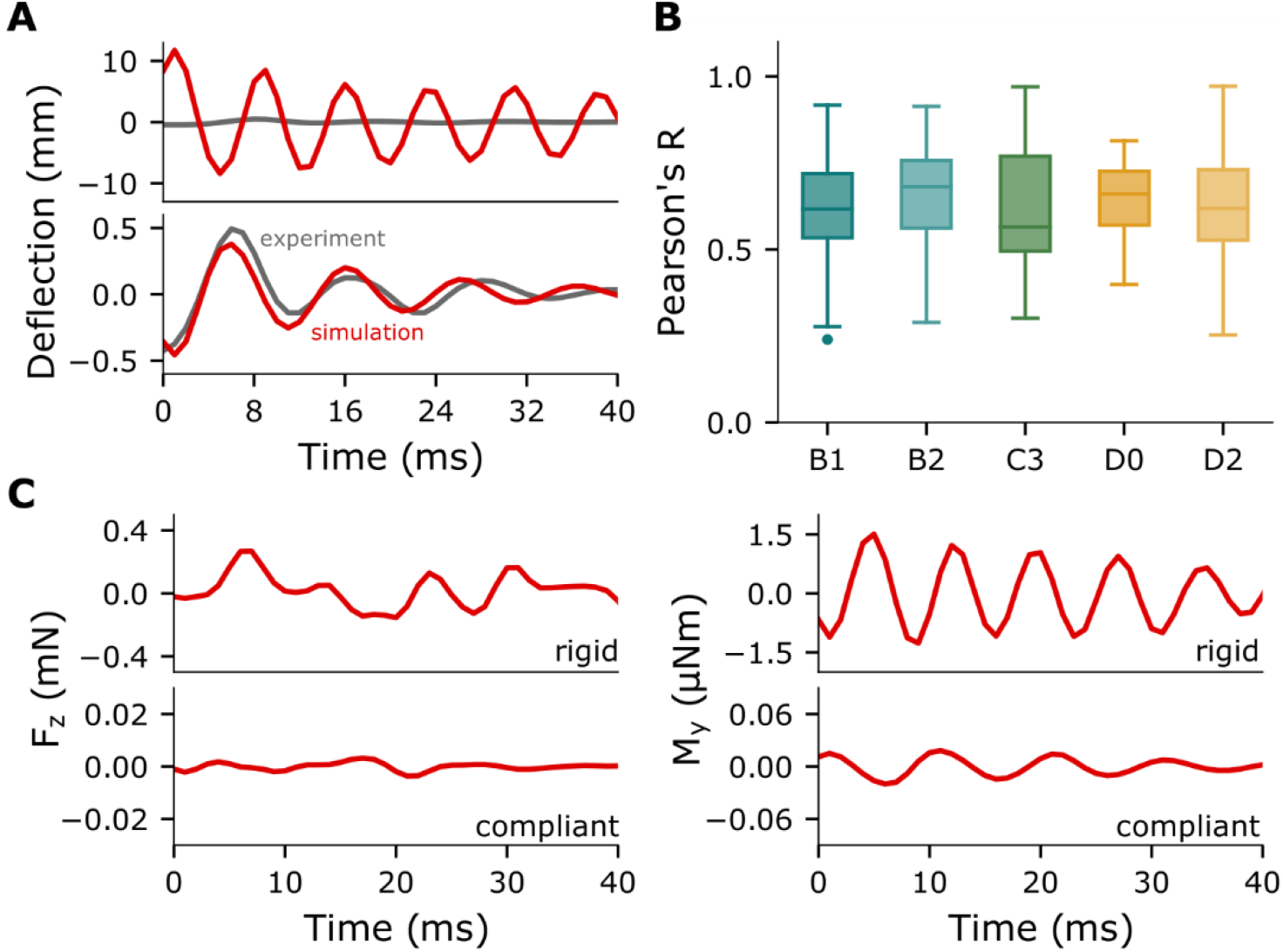
Mechanical properties of the follicle significantly affect whisker dynamics and mechanical signals at the whisker base. **(A)** Displacement of the whisker tip of a representative trial in which the B1 whisker was deflected. The grey traces in both panels show the oscillations measured experimentally, which include the compliant properties of the follicle. The red traces show simulation results when assuming a rigid follicle condition for experiment (upper panel) and after incorporating the compliant properties of the follicle (lower panel). **(B)** Pearson correlation between experimental and simulated trajectory of the most distal link for five different whiskers, labeled on the x-axis, pooled over horizontal and vertical directions. Distributions were computed across all trials for each whisker. R = 0.61 (±0.14) for B1 whisker, R = 0.66 (±0.15) for B2 whisker, R = 0.61 (±0.19) for C3, R = 0.63 (±0.12) for D0 whisker, and R = 0.61 (±0.12) for D2 whisker **(C)** Force in bending direction, *F_z_* (left), and moment about the bending axis, *M_y_* (right) in rigid (upper panel) and compliant (lower panel) follicle condition.

As shown in Figure 4B, the Pearson correlation coefficients between experiment and simulation for both horizontal and vertical deflections for each whisker achieve an average R value of 0.63 (±0.15). These results indicate that the model can robustly predict dynamics at the whisker base even after accounting for the compliant properties of the follicle embedded within the tissue

As expected, the difference between rigid and compliant follicle is also evident in the mechanical signals predicted to occur at the whisker base. Figure 4C shows an example of the forces in bending direction (*F_z_*) and moment about the axis of rotation (*M_y_*) for a B1 whisker corresponding to the tip trajectory in Figure 4A. Compared to the rigid follicle model, the compliant follicle model reduces *F_z_* and *M_y_* by more than an order of magnitude while it also exhibits low pass properties smoothing the mechanical response.

### 2.5. Mechanical signals across the full rat vibrissal array

The organized morphology of the rat vibrissal array plays an important role in shaping tactile input (23, 24, 45, 46). WHISKiT incorporates this morphology (51) to permit simulation of the complete mechanosensory input to the system during naturalistic behaviors.

Figure 5 compares the tactile signals of the rat vibrissal array for three scenarios. In scenario 1 (Figure 5A), a vertical peg is simulated to brush through the center of the immobile array. The peg moves at constant speed (0.4 m/s) from rostral to caudal. Scenario 2 (Figure 5B) simulates active bilateral whisking against two fixed, vertical pegs. Each whisker is driven at its base according to established kinematic equations for whisking motion (29). One cycle of protraction and retraction of the array lasts 125 ms, equivalent to a whisking frequency of 8 Hz. The pegs are positioned laterally, 30 mm from the origin of the array (base point average) with an offset of 10 mm from the nose tip. Finally, in scenario 3 (Figure 5C), the whiskers perform the identical whisking motion as in scenario 2, but the array is positioned in front of the opening of a drainpipe, so that the rat is simulated to actively palpate a typical object found in a natural rat habitat (Figure 5C). Supplementary videos SV1, SV2, and SV3 show the simulations of scenarios 1, 2, and 3, respectively.

**Figure 5.**
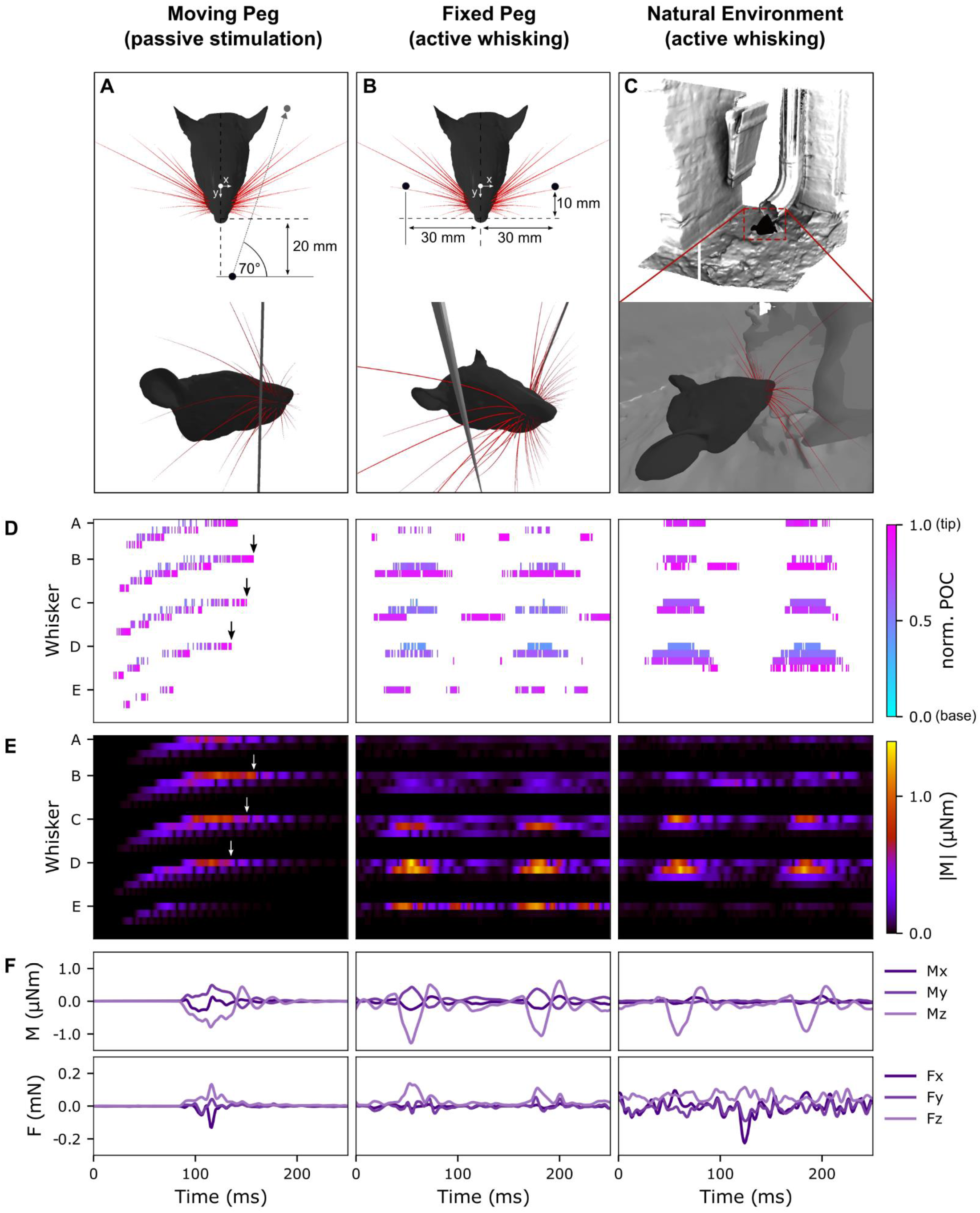
Mechanical response of the full rat whisker array for three scenarios (PP, PA, NEA). For simplicity, only results of the right side of the array are shown. **(A)** Visualization of the passive stimulation experiment. A vertical peg was simulated to move from rostral to caudal through the middle of the immobile right array. **(B)** Visualization of active whisking against two vertical pegs. The array performed a typical whisking motion as described in (29) with a whisking frequency of 8 Hz. **(C)** Visualization of NEA experiment. Same as **(B)**, but array makes contact with the shape of a drainage pipe representing natural environment. **(D)** Points of contact (POCs) for each whisker of the right array over time for each scenario. The POC is normalized to the length of the whisker and indicated by the colormap. The whisker identities are sorted by column (caudal to rostral) and by row (ventral to dorsal). Columns 0 refers to the most caudal column, the Greek column, whereas row 1 refers to most dorsal row. **(E)** Magnitude of base moment (M) of each whisker indicated by color. The sorting of the whiskers is consistent with (D). **(F)** Example of all 6 signal components at the base of the δ whisker for each scenario. All panels share the same time scale (x axis).

Figure 5D shows the point of collision (*POC*) of each whisker in the right array as a function of time for each of the three scenarios. The *POC* is normalized between 0 (whisker base) and 1 (whisker tip). For passive stimulation (left panel), the spatial arrangement and geometry of the whiskers result in a systematic pattern of whisker activation. The moving peg first collides with the most rostral whiskers of the C, D, and E rows. Contact durations are short because the whiskers are small. As expected, contact durations increase as the peg makes contact with the more caudal and larger whiskers. Note that the slip-off of each whisker tip is clearly visible, as reflected in high *POC* values at the very end of each whisker contact.

In contrast, contact responses during active whisking (Figure 5D, middle panel) are more clustered within the whisk cycle. The morphology of the array allows the longer whiskers to touch the pegs, while many of the short, rostral whiskers (columns 3-5) show no contact at all. Because the pegs have a small diameter and are oriented orthogonal to the whisking motion, some whiskers slip past the pegs and cause secondary collisions during retraction. This effect is particularly observable for the A2, C2, D2, and E1 whiskers. Interestingly, with an extended surface such as the drainpipe (Figure 5D, right panel), this type of slip-off is less likely, as seen in the highly clustered contact patterns during protraction.

Figure 5E shows the magnitudes of the bending moment, *M*, at the base of all whiskers during each behavioral condition. In the left panel, oscillations in the bending moment reflect each whisker’s vibrations after it has slipped off the peg (black and white arrows). This effect is particularly visible for whiskers β, γ, and δ. Although less distinct, similar vibrations also cause *M* to increase between whisks during both active whisking scenarios (Figure 5E, middle and right panels). Also note *M* and *POC* tend to be inversely related, because contact closer to the whisker base generally results in a greater mechanical response. Finally, because the head was stationary across all simulations, the more caudal whiskers always undergo larger deflections, leading to increased values of *M* and increased probability of lower *POC* in this region of the array.

An example of the individual signal components at the base of a single whisker (in this case, δ) is shown in Figure 5F. For this whisker, the bending moment during active whisking appears qualitatively similar between the peg and drainpipe conditions. However, these two conditions are clearly distinguishable at the level of the entire array (Figure 5E).

## 3. Discussion

Mechanical responses of isolated whiskers have been quantified in various contexts (33, 40, 43, 47–53), but models of the vibrissal system have typically been quasi-static, two-dimensional (2D), and limited to a single whisker (32, 34, 38, 42, 44, 54–60). These are significant limitations when studying neural circuitry that has evolved with multiple sensors interacting with a three-dimensional complex world. Behavioral studies have shown that natural rodent whisking behavior involves contact and collisions of many whiskers with the environment. Studies of the barrel cortex indicate that neural information is spatially integrated across multiple whiskers and that the statistics of the stimuli have a significant effect on the receptive fields due to adaption mechanisms (61). These findings indicate that it is crucial to study neural activity in the vibrissal system in the context of natural behavior and the full whisker array.

The 3D dynamical model of the rat vibrissal array developed here contains simplifying assumptions. First, the individual whisker is modeled as a chain of 20 rigid links connected with torsional springs, each with two degrees of freedom. Because the spatial resolution of the whisker is limited to 20 data points (links), some dynamic behavior such as higher order resonance modes cannot be simulated. These modes are presumed to play a role in texture discrimination (34, 39, 62), but they lie beyond the scope of the present model. Second, the model assumes homogeneous Young’s modulus and damping properties within and across whiskers. However, previous work has suggested that Young’s modulus varies along the whisker, especially due to the presence of the medulla (the hollow portion of the whisker), and the cuticle (35). Differences in density between proximal and distal whisker regions have also been shown; this variation was approximated as a linear density increase from base to tip (63).Third, because each node is limited to two degrees of freedom, shear forces cannot be modeled. Although beam theory suggests that shear forces in straight vibrating beams with an aspect ratio greater than 100 (aspect ratios of whiskers are 1000 or more) are negligible (64), it remains unclear how taper and curvature of the whisker affect the shear forces at the base and if and how they are transduced by the mechanoreceptors embedded in the follicle.

The present model was not developed to replicate the sensory input of a single, individual rat at high precision. Differences in animal size, age, sex, and strain ultimately manifest in different sizes, scales, shapes, material properties and spatial arrangement of the vibrissae. Despite this variability, rat brains have found solutions that allow these animals to use their vibrissal systems to accomplish similar tasks. Understanding the underlying principles of the rat somatosensory pathway therefore cannot rely on modeling the sensors of an individual rat in detail (65). Although existing single-whisker models may be able to predict the dynamics of individual whiskers more accurately (32, 38, 51), the present model has been validated to generate reasonable approximations of real whisker dynamics, well within the naturally occurring variability within and across animals. The present model can be viewed as a “prototypical” rat, whose parameters and dynamics fall within the range of biological rats.

We anticipate that *WHISKiT Physics* will open new ways to test hypotheses about the structure of mechanical signals across the rat whisker array; these data are currently inaccessible in the real animal. We also expect it to enable the field to leverage techniques such as information theory, virtual reality, and reinforcement and machine learning. These approaches will allow researchers to model both sensory information processing as well as control circuits, learning, and environmental interactions. Finally, we anticipate that the simulation system can be used to bootstrap hardware implementation of artificial whisker systems, which could potentially accelerate and facilitate the design process of whisker-based robots.(56, 57, 60, 66, 67).

## 4. Methods

All experiments involving animals were approved in advance by the Animal Care and Use Committee of Northwestern University.

### 4.1. Model of a single whisker

The model of a single whisker was created using the Bullet Physics Library (68) and extends a previous two-dimensional (2D) model (51) to three dimensions (3D). The whisker is modeled as a chain of N conical frustums (links) connected by N-1 equidistant joints (nodes). The length of the links *l_link_ = S/N* depends on the whisker arclength S and the number of links N, while the radius at each node *r_n_* decreases linearly from the whisker base to the tip (eq.1). Schematics of two views of the model are shown in Figure 6.

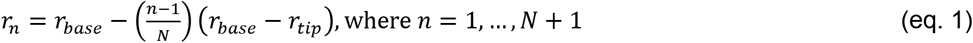

**Figure 6.**
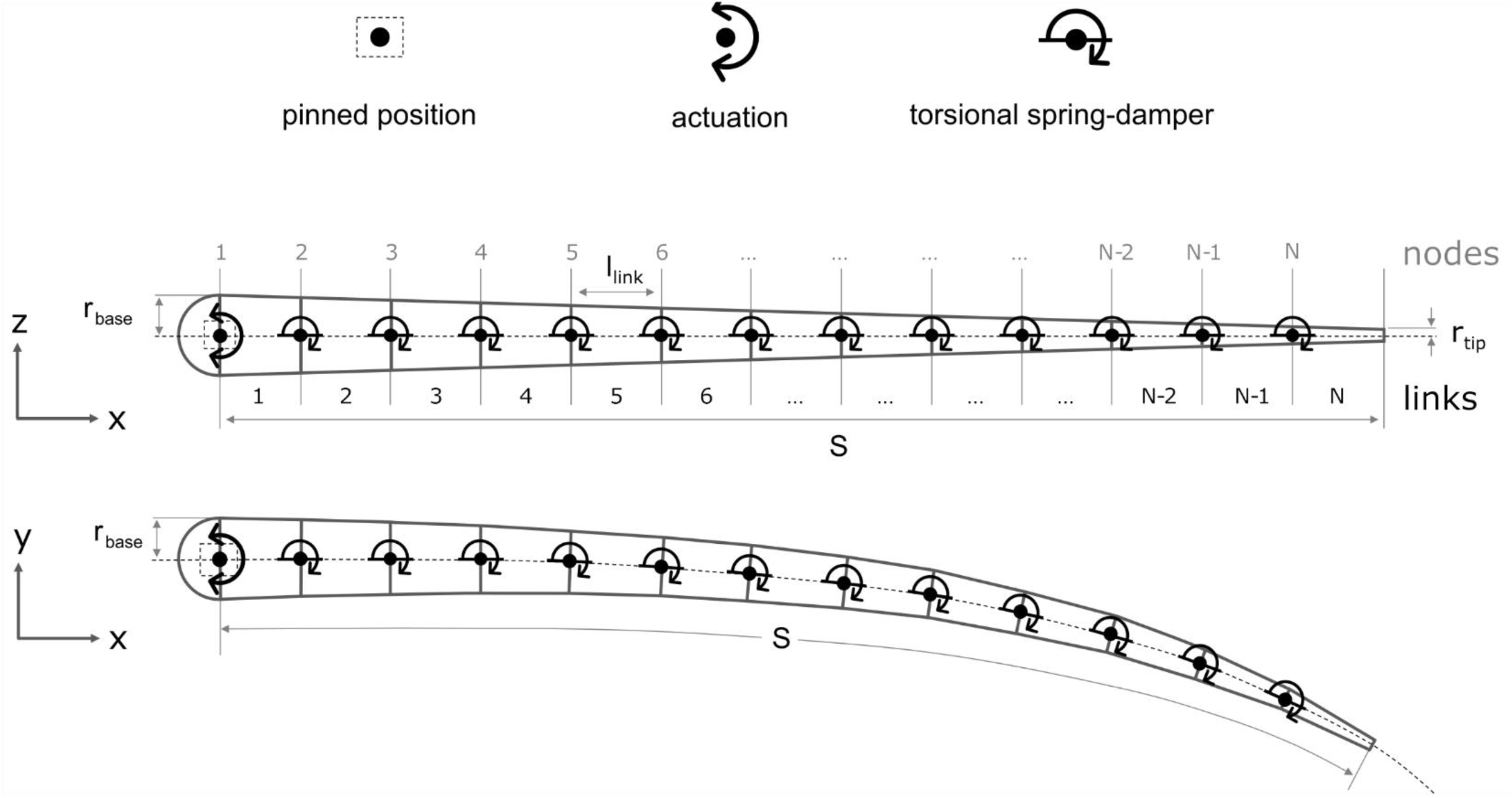
The schematics illustrate the whisker model used in all simulations. The whisker is rigidly driven from node 1, which represents the follicle and is pinned so that it can rotate but not translate. Nodes 2 – N each represent a 3D torsional spring damper. The whisker is straight in the x-z plane and has intrinsic curvature in the x-y plane. In each simulation the curvature was chosen to be appropriate for the specific whisker being simulated (see text for details).

Each link has mass 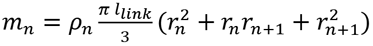, where density 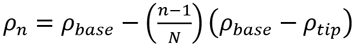 increases linearly from base to tip. Values for density at the base (*ρ*_*base*_) and the tip (*ρ*_*tip*_) were obtained from a previous study (63).

Nodes 2 through N are modeled as torsional spring-dampers with two degrees of freedom, permitting rotations about the y and z axes. The parameters *k*_*n*_ and *c*_*n*_ represent stiffness of the spring and damping in the two bending directions (rotations about the y and z axes). Twist of the whisker (rotation about the x-axis) was omitted. The whisker follicle is represented by node 1. The follicle is pinned (cannot translate) and its rotation about all three axes is rigidly controlled. A torsional spring-damper was later added to model the tissue elasticity of the follicle in the skin (section 2.4), allowing rotations about the y and z axes.

Theoretically, each node can be viewed as the pivot point of a pendulum (51). The adjacent distal portion of the whisker has mass *M*_*n*_, which determines the mass of the pendulum, and the distance *L*_*com,n*_ between its center of mass (com) and the n^th^ node determines the length of the pendulum. Thus, the stiffness and damping of the spring associated with the node can be calculated as:

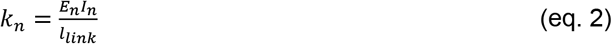

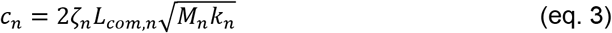

where *E*_*n*_ is Youngs’s modulus and *I*_*n*_ is the area moment of inertia of the associated link. The variable *ζ*_*n*_ represents the damping ratio of the spring of the n^th^ node.

From equations 2 and 3 and 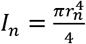, it is clear that the stiffness and damping at each node are primarily determined by the geometry of the whisker. Following the methods of previous studies (32–35, 38, 50, 51, 69), our model assumes uniform Young’s modulus and uniform damping ratio, i.e., *E*_0_ = *E*_1_ = ⋯ = *E*_*n*_ and ζ_0_ = ζ_1_ = ⋯ = ζ_*n*_. We denote the corresponding model parameters as *θ*_*E*_ and *θ*_*ζ*_. We optimized parameters *θ*_*E*_ and *θ*_*ζ*_ using kinematic data obtained from a real whisker. The experimental setup and the optimization method are explained in section 4.3.

### 4.2. Temporal resolution and spatial accuracy of the model

The temporal resolution and spatial accuracy of the whisker model depend on three factors: *1)* the sampling frequency of the physics simulation; *2)* the number of iterations of the physics solver; and *3)* the number of links used to model the whisker. Increasing the sampling frequency and number of iterations and links increases both temporal and spatial accuracy but also increases computational load and simulation time.

Previous studies have measured first mode fixed-free resonance frequencies up to 360 Hz (C4 whisker) in response to pulse stimulation and up to 585 Hz in response to a piezoelectric chirp stimulus (34). Our model is primarily intended to simulate vibrations resulting from pulse-like stimuli similar to collisions. Thus, we limit the temporal resolution of the model to resonant modes less than 500 Hz with a sampling frequency of 1kHz (1 ms per time step).

The spatial accuracy of the whisker dynamics depends on the number of links of the whisker model and the number of solver iterations. The number of links is a model design choice that affects the flexibility of the whisker as well as the computation time, which is proportional to the number of simulated bodies. A battery of initial simulations showed that a model with 20 links and 40 solver iterations was a good compromise between accuracy and efficiency.

### 4.3. Experiments for model optimization and model validation

To optimize and validate the model we performed two separate experiments. The first experiment involved rotating whiskers on a motor in the horizontal plane, resulting in motion that was approximately 2D although 3D motion was quantified. Data from this first experiment were used to optimize material parameters (Young’s modulus and damping coefficient) for the model. The second experiment involved manual deflections of different whiskers in the anesthetized animal. The whiskers were deflected in several different directions, and these data were used to optimize the constants of the spring modeling the elasticity of the follicle.

The first experiment used two whiskers, α and B1, whose geometric parameters are listed in Table 2. Both whiskers were trimmed so that their tips could be clearly seen in the video, therefore they both have a shorter arc length and larger tip diameter than typical. The base of each whisker was fixed to the vertical shaft of a DC motor and rotated in a “gaussian pulse” motion. The whisker was oriented so that its intrinsic curvature coincided approximately with the horizontal plane. The motor was controlled using a microcontroller (PIC32) running a feedback controller at 5 kHz. At the beginning of each trial, the initial motor position was set to 0° and the microcontroller was synchronized with two orthogonally-mounted high-speed video cameras (Mikrotron 4CXP; E1: 1000fps, E2: 500 fps) used to track the whisker’s motion. The amplitude, speed and frequency of the driving signal was varied (Table 2) and each parameter combination was repeated five times. After data collection, the whisker was removed from the motor and the base diameter (at the fixation point on the motor) and length were measured using a microscope (Leica DM750).

**Table 2.** Dimensions of the whiskers and motor parameters used in Experiment 1.

| Dimensions of the whiskers |  |  |  |
| --- | --- | --- | --- |
| Whisker | Arc length (mm) | Base diameter ($\mu\text{m}$) | Tip diameter ($\mu\text{m}$) |
| $\alpha$ | 29.6 | 120 | 35 |
| B1 | 34.1 | 131 | 30 |
| Motor parameters |  |  |  |
| Parameter | Variable | Values |  |
| Driving signal | N/A | $\varphi(A, \omega) = Ae^{-\omega(t-t_0)^2}$ | |
| Amplitude (deg) | A | 45 |  |
| Speed (deg/s) | $\omega$ | 900, 2000, 3500 | |
| Frequency (Hz) | f | - |  |
| Time offset (s) | $t_0$ | 45/200 | |

The second set of experiments was performed in the anesthetized rat as part of a separate study (Bush et al. 2019, in review). All whiskers except for one were trimmed down to the length of the fur. The spared whisker was manually deflected with a graphite probe in eight cardinal directions, at two or three different contact points along its length, at two different speeds.

In both Experiments 1 and 2, the 3D whisker reconstruction from the two orthogonal camera views was performed in three steps. First, the whiskers were tracked in 2D using the software “Whisk” (70). Tracking was manually verified frame by frame in both views. In each view, the 2D tracked whisker shapes were cleaned by first removing and interpolating over mistracked basepoints via a median filter (window = 5 frames) and outlier deletion (Grubbs test *α* < 1^−8^), and then entire 2D whisker shapes were smoothed with a spatial LOWESS filter (span=15% of whisker length). Second, the two cameras were calibrated with the Caltech Camera Calibration Toolbox, OpenCV, and custom Matlab and python code. Finally, an iterative optimization was used to find the best 3D whisker shape that minimized the 2D back-projection error, defined as the Euclidean distance between the back-projected whisker and the actual, imaged whisker, summed over all back-projected points. Fit quality was inspected manually by observing the shape of the 3D whisker over time, viewing the overlap of the 2D back-projections with the original tracked 2D whiskers, and by monitoring the temporal trajectories of the base and tip points for large deviations.

To analyze the manual deflections in Experiment 2, we used semi-automated tracking code to determine the 3D contact point and time at which the graphite probe made contact with the whisker. Tracking quality in all videos was monitored online during tracking and confirmed as accurate offline after tracking by a second user.

The final 3D tracked data occasionally exhibited large errors due to tracking problems in the 2D data, the 3D reconstruction process, or calculation of the contact point of the probe. Such errors, which were easily observed as exceptionally large outlier “glitches,” considerably decreased the correlation measured between experiment and simulation. Therefore, the 3D tracking data was filtered with a thresholded median (Hampel) filter using a window size of 3 frames and a threshold of 1.5 standard deviations for outliers. In Experiment 2, data were upsampled from 500 fps to 1000 fps to match the sampling rate of the simulations.

Finally, the best trials were manually selected for each whisker in both experiments. The resulting datasets are shown in Table 3.

**Table 3.** Datasets for Experiment 1 and Experiment 2.

|  | Whisker identities | Number of trials |
| --- | --- | --- |
| Experiment 1 | $\alpha$ | 10 |
|  | B1 | 6 |
| Experiment 2 | B1 | 83 |
|  | B2 | 63 |
|  | C3 | 13 |
|  | D0 | 28 |
|  | D2 | 74 |

### 4.4. Model optimization

Two optimizations were performed to match the simulated dynamics with the dynamics of real whiskers. The first optimization determined optimal values of the model parameters of the single whisker model (*θ*_*E*_ and *θ*_*ζ*_) using data collected *ex vivo* (experiment 1), while the second optimization was used to find the optimal values for the follicle parameters (stiffness and damping) using data collected *in vivo* (experiment 2).

#### 4.4.1 Optimization of model parameters *θ*_*E*_ and *θ*_*ζ*_

In order to optimize the parameters *θ*_*E*_ and *θ*_*ζ*_, the motor setup described for experiment 1 was replicated in simulation using Bullet Physics Library (68, 71).

To reduce the computational load of the simulations, the 3D-merged data was down-sampled from 64 to 20 nodes by linearly interpolating the coordinates of the whisker to 20 equidistant points between the axis of rotation and the tip of the whisker at each time step. The motor driving signal of was smoothed to remove high frequency discontinuities associated with the quantization of the motor encoder.

Because whisker dynamics depended on the geometry of the whisker, the single whisker model was modified to accurately match the dimensions and shape of the real whisker in the experiment (Table 2). The 3D data points of the first frame were used to reconstruct the whisker in simulation. These data points defined the locations of the nodes and the length of the links between them, respectively. The node at the base was placed at the origin (0, 0, 0) of the world frame. The smoothed driving signal of the motor was used to control the angular displacement of the whisker base about the axis of rotation.

We used a brute-force approach to iterate through 1296 combinations of *θ*_*E*_ and *θ*_*ζ*_ values within a specific range (*θ*_*E*_ was varied between 2.0 and 6.5 GPa and *θ*_*ζ*_ between 0.15 and 0.6). For each pair of parameter values, five trials were randomly sampled for each of the two whiskers. Then each trial was simulated and evaluated in terms of its first-mode resonant frequency (FRF), logarithmic decrement δ, and peak amplitude A, computed from the y trajectory of the whisker tip evolving over 1000 samples (= 1 second). The FRF was determined by finding the peak of the power spectrum computed via the Fast Fourier Transform (FFT). The value of δ was calculated in the time domain. Given the magnitudes of the first two adjacent peaks, y_0_ and y_1_, 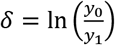. The value of y_0_ was used as measurement for peak amplitude. We used the median symmetric accuracy (MSA) (72) to quantify the total error of the simulations across the 10 trials. The MSA was computed using (eq. 4a), where simulated and experimental measurements are denoted by the subscript _sim_ and _exp_, respectively. The median was calculated from the pooled FRF, δ, and A measurements across all 10 trials.

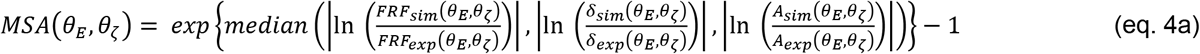

The optimal parameter values 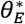 and 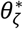 were obtained by finding the parameter combination yielding the minimum MSA across all parameter evaluation.

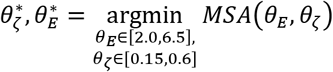

Across all trials used during optimization, high Pearson correlation coefficients for both whiskers, α (x: 0.92, y: 0.92) and B1 (x: 0.99, y: 0.78), indicate a close match between simulation and experiment for both whiskers.

#### 4.4.2. Optimization of the follicle parameters

To determine the parameters that describe the mechanics of the follicle embedded in the skin, we used data from Experiment 2. The follicle was modeled as a torsional spring-damper system with two degrees of freedom, allowing rotation about the y and z axes. In contrast to the single whisker model, the stiffness and damping parameters of the torsional spring-damper system in the follicle are not directly related to any specific material properties.

Given that the oscillations of the whiskers in Experiment 2 were very small, the logarithmic decrement δ was difficult to measure. The error metric in eq. 4a was therefore adjusted to include only the error in FRF and the error in the magnitude of the first two adjacent peaks y_0_ and y_1_ of the whisker tip oscillations (eq. 4b).

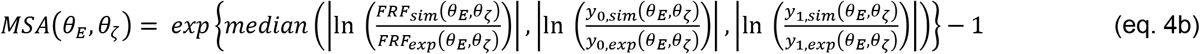

We then followed similar procedures as for the optimization of model parameters *θ*_*E*_ and *θ*_*ζ*_. Whiskers B1, B2, and D2 from Experiment 2 were used. For each parameter combination, three trials were randomly sampled for each whisker and the MSA computed according to eq. 4b. Again, the optimal values were determined by finding the minimum error across all simulated parameter combinations.

### 4.5. Validation experiments

#### 4.5.1. Extrapolation of optimized material parameters across whiskers with varying geometries

After optimizing and validating dynamics for a select number of whiskers in experiment 1, we validated the resonance behavior of all whiskers across the array. To perform this validation, we searched the literature for studies that had published values of whisker resonance along with whisker length, base diameter, and tip diameter. We could find no study that had published all of these values for the same set of whiskers. However, Hartmann et al. (33) published resonance frequencies, arc length, and base diameter for 24 whiskers, and we were able to obtain tip diameter measurements for the same 24 whiskers from the original 2003 experiment (supplementary Table 1).

Based on these data, we simulated 24 straight whiskers with the measured arc lengths and base and tip diameters. The material parameters were set to the optimized values *θ*_*E*_ = 5.0 and *θ*_*ζ*_ = 0.33. In simulation, the base of each whisker was rigidly clamped to a motor (defined as the origin) such that the whisker lay in the horizontal (x-y) plane and was aligned with the x-axis. The motor rotated the whisker about the vertical (z) axis. Each simulation lasted 1 second, sampled at 1kHz. The angle from the original position ν(t) was changed according to a Gaussian-like function with 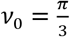, μ = 4σ and σ = 0.025 to ensure smooth motor movement at the beginning of the actuation according to eq. 5:

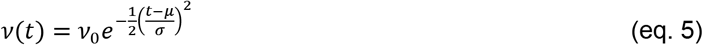

The rotation of the motor was terminated abruptly after 0.128 seconds to induce vibrations of the whisker. The simulation was performed once for each whisker geometry.

The x, y, and z position of all 20 links was recorded at each time step, but only the y-component of the whisker tip (link 20) was used for further analysis. The first 125 samples were removed from each trial so that the data contained only the free oscillations of the whisker. An FFT was performed on the remaining 875 samples, and the frequency with maximum amplitude was found to identify the first resonant mode.

#### 4.5.2. Extrapolation of optimized material parameters to the average rat vibrissal array

The rat vibrissal array consists of 31 whiskers on each side, a total of 62, neatly organized in a grid of approximately 5 rows and 7 columns. Each location is annotated by the associated row (letters A-E from dorsal to ventral) and column (numbers 1-6 from caudal to rostral). The whiskers of the first (most caudal) column, which is slightly offset in respect to the rest of the array, are denoted by Greek letters (α, β, γ, δ from dorsal to ventral).

The arc length (S), base radius (r_base_), and the radius slope of the whisker can be approximated as functions of whisker position (row, column) from which the tip radius can be calculated. The intrinsic curvature of the whisker is approximated by a quadratic function *y* = *Ax*^2^ in the x-y plane according to Belli et al., where *A* represents the curvature coefficient (Belli *et al.*, 2018). The value for A can be calculated by using the relationship between intrinsic curvature and arc length of the whisker. In our model, N equidistant points along this quadratic curve represent the position of the nodes.

We tested (Figure 2B, lower panel) whether the simulations using optimized parameters would generalize to explain the dynamics of all whiskers within the average whisker array as described by the equations found in Belli et al. (40). The simulations identical to those described in section 4.5.1, but all 28 whiskers comprising the right side of the array (rows A-E, columns Greek-5) were simulated.

The experimental data for comparison (resonance frequency and arc length) was compiled from Hartmann et al. (33), Wolfe et al. (39) and Neimark et al. (34), see supplementary Table 1. The data from Wolfe et al. (39) were obtained from Figure 4 of the article. The resonance frequencies from Neimark et al. (34) was obtained from Table 1 of the article (Pulse mode experiment). Because whisker arc length was not published for these data, we used average values from Belli et al. (40) for the corresponding whisker identities (β, C1, C2, C3, C4).

#### 4.5.3. Collision with a straight edge

To validate simulation dynamics during collision, we compared results with a previously published analytical model (32) that simulated a straight whisker colliding with the edge of an object. For this purpose, each simulated whisker was rotated at its base at constant angular velocity about the vertical axis through the base point until the 19th link (95% of whisker length from the base) collided with a rectangular edge. Initial angular displacement from the edge was set to 5.6° 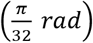. When contact with the object was detected, the driving force at the base was set to zero, allowing natural deceleration of the whisker. Collision was set to be inelastic to avoid rebound.

The collision angle between whisker and edge was defined to be 60° in the horizontal, and 45° in the vertical plane to generate mechanical responses in all three dimensions. We performed simulations with 12 whiskers from 4 different rows (α, A2, A2, B1, β, B1, B2, γ, C1, C2, δ, D1, D2). In the first set of simulations, intrinsic curvature was omitted, i.e., 12 straight whiskers replicated the experiment described in Boubenec et al. (32). In the second set of simulations, all whiskers were curved according to the model in Belli et al. (41). For each whisker, we simulated collisions for 6 velocities (90, 180, 270, 360, 450, 540 deg/s) while recording all six components of the mechanical signals at the base (*Fx*, *Fy*, *Fz*, *Mx*, *My*, *Mz*) as well as a binary “contact vector,” *C*, which indicated contact state (0 or 1) for each whisker link. Simulation output was sampled at 1 kHz and simulations were terminated after 0.2 seconds.

For signal analysis, the mechanical components *Fx, Fy, Fz, Mx, My, Mz* were smoothed by convolving a 10^th^ order Hanning window. Time of collision was obtained by finding the first nonzero value of *C* of any whisker link.

To determine the magnitude of the shock immediately after collision for each whisker, the derivative of the signal Mz was rectified and the maximum *Ṁz_max_* at time *t_max_* within the first 10 ms after collision was found. The shock magnitude *Ṁz_max_* was normalized for each whisker by dividing by the average across all six speeds. Bending moment and transverse force at time *t*_*max*_ were computed using eq. 6–7.

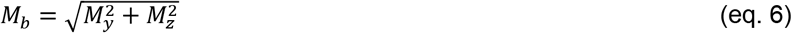

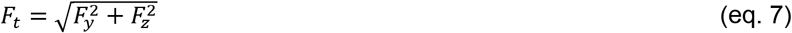

Propagation of the shock wave from tip to base was determined for the two longest whiskers in each of the four rows (α, β, γ, δ, A1, B1, C1, D1). Whiskers A2, B2, C2, D2 were too small and stiff to exhibit a measurable shock wave. To track the propagation of the shock wave, the location of maximum deflection point of each whisker relative to whisker length was measured in respect to position of first contact for 5 subsequent time samples.

### 4.6. Simulation of complete vibrissotactile input across the array

The mechanical signals at the base of each whisker of the full rat whisker array were simulated for three scenarios: motion of a vertical peg through the immobile array, active whisking against two fixed pegs, and active whisking against the 3D shape of a drainpipe, representing the natural environment. In all three scenarios, the whiskers were arranged according to the morphology of the rat whisker array described in Belli et al. (41). For visualization purposes, a scanned rat head was used to model the head to which the array is attached. In all simulations, collisions between the head and other objects were suppressed to increase the simulation speed.

To simulate the first scenario (passive stimulation with the peg), the origin of the whisker array (base point average) was placed at the origin of the world frame and oriented such that the average row plane was approximately parallel to the horizontal plane. The whiskers remained at resting position while a vertical peg moved with constant speed (0.4 m/s) from rostral to caudal through the middle of the array. The peg had a diameter of 1 mm and a length of 80 mm. An illustration of the simulation experiment is given in Figure 5A and supplementary video SV1.

To simulate active whisking against the two pegs, the origin of the whisker array (base point average) was placed at the origin of the world frame and oriented such that the average row plane was approximately parallel to the horizontal plane. Two vertical pegs were placed bilaterally with an offset of ±30 mm in the x axis and −10mm in the y axis to cause collisions with the protracted whiskers. Position and orientation of the array and the pegs remains the same while the whiskers perform a typical whisking motion, as described below. An illustration of the simulation experiment is given in Figure 5B and supplementary video SV2.

Finally, simulations of whisking against a drainpipe serve as an example of how the model of the complete vibrissal array can be used to analyze tactile information acquired in natural environment. In these simulations, the position and orientation of the rat model was manually selected to ensure sufficient but not extreme contact between the surface of the 3D scan and the whisker array. The positions of the rat head and the 3D surface were held fixed while the whiskers performed a typical whisking motion. To model the natural environment of a rat, we collected 3D representations of a typical rat habitat (drainpipe) in the Evanston, IL area with a KINECT™ for Xbox V2 (Microsoft) and a Predator Helios 300 Laptop (Acer). The scans collected covered a volume of approximately 2 cubic meters. In Geomagic® Design X™ (3D Systems, Inc.), the point cloud data was manually edited to remove holes and erroneous points before triangulation to generate a mesh with maximum edge length of 3mm. The final mesh was exported as a Wavefront OBJ file which was imported to the physics simulation. An illustration of the simulation experiment is given in Figure 5C and supplementary video SV3.

For both active whisking scenarios, the whiskers performed sinusoidal whisking motion at a frequency of 8 Hz with a maximum protraction angle of 35 degrees and maximum retraction angle of 15 degrees from rest. Previous work (Knutsen, Biess and Ahissar, 2008) found that natural whisking behavior involves elevation φ(*t*) and torsion ζ(*t*) of the whisker, both of which show a row-wise dependency on the angle of protraction *θ*(*t*). Based on these findings (Knutsen, Biess and Ahissar, 2008), we constructed equations of motion (eq. 8–9) for each row (A-E) which are used to drive angular rotation at the base point during active whisking.

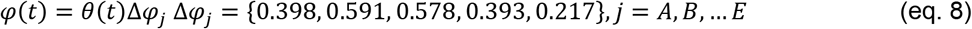

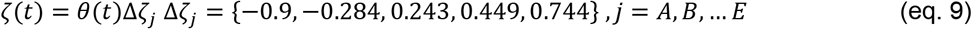

Each simulation lasted 250 ms involving 2 whisk cycles and was sampled at 1kHz. All six components of the mechanical signals at the base of each whisker (*Fx*, *Fy*, *Fz*, *Mx*, *My*, *Mz*) and a binary vector *C* indicating contact (1) or no contact (0) for each whisker link were recorded. For signal analysis, the mechanical components *Fx*, *Fy*, *Fz*, *Mx*, *My*, *Mz* were smoothed by convolving a 10^th^ order Hanning window. The base moment magnitude *M* was computed using eq. 10.

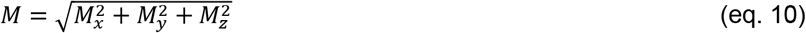

Point of contact (*POC*) was determined by the number of the links in contact relative to total number of links (base: *POC* = 0.0, tip: *POC* = 1.0).

## Notes

#### Summary of Updates

Figures revised.

https://github.com/SeNSE-lab/whiskitphysics

